# Chromatin Patterns Distinguish Breast Tumor Subtypes and Disease Progression in Association with *ANP32E* levels

**DOI:** 10.1101/2021.07.28.454151

**Authors:** Garrett L. Ruff, Kristin E. Murphy, Paula M. Vertino, Patrick J. Murphy

**Author notes:** correspondence &, 601 Elmwood Avenue, Rochester, NY 14624.

## Abstract

Despite highly advanced diagnosis and treatment strategies, breast cancer patient outcomes vary extensively, even among individuals with the same diagnosis. Thus, a better understanding of the unique molecular characteristics that underlie tumor trajectories and responses to therapy remains a central goal. We report that chromatin patterns represent an important characteristic, capable of stratifying tumor identity and progression. We find that patterns of chromatin accessibility can be classified into 3 major groups, representing Basal-like tumors, hormone receptor (HR)-expressing tumors, and invasive lobular Luminal-A tumors. Major chromatin differences occur throughout the genome at motifs for the transcription factor FOXA1 in HR-positive tumors, and motifs for SOX9 in Basal-like tumors. A large portion of lobular Luminal-A tumors display a chromatin signature defined by accessibility at FOXA1 binding motifs, distinguishing them from others within this subtype. Expression of the histone chaperone *ANP32E* is inversely correlated with tumor progression and chromatin accessibility at FOXA1 binding sites. Tumors with high levels of ANP32E exhibit an immune response and proliferative gene expression signature, whereas tumors with low ANP32E levels appear programmed for differentiation. Our results indicate that ANP32E may function through chromatin state regulation to control breast cancer differentiation and tumor plasticity.

## INTRODUCTION

Cellular programming is controlled by epigenetic modifications, transcription factor binding patterns, and DNA packaging within the nucleus. These mechanisms control how gene transcription machinery gains access to DNA at transcription start sites (TSS) and cis-regulatory enhancers, ultimately controlling cellular programming through regulation of gene expression. Regions with more accessible chromatin tend to be more highly transcribed, and inaccessible regions are typically silent (1). Overall, chromatin accessibility is generally stable in terminally differentiated cells, along with steady gene expression profiles, and the majority of chromatin state dynamics occur either during embryonic development or as a consequence of disease progression, including during carcinogenesis (1–3). Breast cancer is among the most frequent and well-studied forms of cancer worldwide, but chromatin state specific differences among breast cancers have not been established.

Breast cancer represents the most diagnosed cancer in women (4) with an estimated 2.1 million newly diagnosed cases globally in 2018 (5). Measurements of gene expression or protein abundance have enabled breast tumors to be classified into discrete subtypes, allowing for diagnosis-specific treatment strategies based on underlying cellular programming. Measurements of chromatin states have the potential to provide additional insights into breast cancer mechanism and may ultimately lead to novel therapy strategies. For example, a recent study of 410 tumors from The Cancer Genome Atlas (TCGA) used chromatin accessibility measurements to identify more than 500,000 putative gene regulatory elements, including thousands of genomic locations where accessibility differences occurred in a disease-specific and tissue-specific manner (2). Separate studies of myeloma have also found that accessibility levels at gene-distal enhancer regions enable accurate prediction of nearby oncogene expression levels, as well as cancer subtype classification (6). Similar breast cancer focused studies are lacking and have the potential to identify parallel associations.

Presently, breast cancer diagnostic methods include histologic classification, which is based on the expression of estrogen receptor (ER) and progesterone receptor (PR), as well as expression or amplification of the *ERBB2*/HER2 (gene/protein). These measurements are critical in selecting patients for hormonal or HER2-directed therapy. Triple-negative breast cancers (TNBC) lack expression of these biomarkers (7), and treatment options are thus limited to chemotherapy. PAM50 (Prediction Analysis of Microarray 50) gene expression profiling is an effective method to identify “intrinsic subtypes”, classified as Luminal A (Lum-A), Luminal B (Lum-B), HER2-enriched, and Basal-like (Basal-L), and these subtypes provide initial insight into molecular mechanisms, likelihood of progression, and patient outcome (8). While much is known about the biological underpinnings of these subtypes, understanding how they correspond with chromatin accessibility differences may uncover additional mechanisms, including novel roles for chromatin and epigenetic factors.

The vast majority of transcription factors can only bind DNA at accessible chromatin locations, rendering them non-functional at inaccessible binding sites (1). When chromatin state changes occur, increased transcription factor binding generally leads to increased expression of neighboring genes (1–3). Many transcription factors which are normally active during development become reactivated in breast cancer, and depending on chromatin state, these factors may influence tumorigenic behavior. For example, SOX9 and FOXC1 are important for developmental regulation of transcription in multipotent neural crest stem cells (9, 10), and they become reactivated in breast cancer to co-regulate Basal-L cancer initiation and proliferation (11). In contrast, FOXA1, which is normally active in hematopoietic progenitor cells, acts coordinately with ER to suppress Basal-L programming and reinforce the luminal phenotype (12, 13). Furthermore, hyperactivity of FOXA1 promotes pro-metastatic transcriptional programs in endocrine-resistant tumors (14, 15). Thus, assessing chromatin accessibility in breast cancer tumors, at specific transcription factor binding sites, could be highly informative for studying the molecular function of numerous factors already known to control cancer outcomes.

We recently defined the histone chaperone protein ANP32E as a genome-wide regulator of chromatin accessibility in mouse fibroblasts (16). ANP32E functions to modulate the installation/removal of H2A.Z from chromatin, regulating chromatin remodeler activity and limiting chromatin accessibility. We found that loss of ANP32E caused thousands of gene promoters and enhancers to become more “open”, leading to activation of neighboring genes. These changes were accompanied by cellular reprogramming events where loss of ANP32E caused cells to take on a more differentiated transcriptome phenotype. Interestingly, a recent study suggests that ANP32E may be an independent prognostic marker for human breast cancers, where higher ANP32E protein levels are associated with the TNBC subtype and correlated with a shorter overall and disease-free survival. Moreover, forced downregulation of ANP32E suppressed TNBC tumor growth in xenograft models (17). However, the precise mechanisms by which ANP32E functions to support breast cancer growth and its role in defining breast cancer phenotypes has not been fully established.

To gain insight into chromatin state function and heterogeneity in human breast cancer, we used an unsupervised computational approach to segregate tumors into defined groups based solely on genome-wide chromatin accessibility patterns. Basal-L tumors segregated as a homogeneous class within group 1, where as a mixture of tumor types was found within group 2, including nearly all Lum-B and HER2-enriched tumors, and group 3 consisted primarily of lobular Lum-A tumors. By defining the chromatin accessibility ‘signature’ associated with each group, we identified DNA sequence motifs for specific transcription factors. SOX9 motifs were most accessible in group 1 tumors, and FOXA1 motifs were most accessible in HR+ tumors within groups 2 and 3. Finally, we found that expression for the chromatin factor ANP32E was anti-correlated with tumor progression and with accessibility at FOXA1 binding sites among group 2 and 3 tumors, suggestive of a novel mechanism by which FOXA1 activity may be regulated in breast cancer tumors. Our results highlight the potential for future disease focused studies of chromatin accessibility, as well as epigenetic therapies directed at disrupting chromatin regulatory factors.

## Materials and Methods

### Measurements of Chromatin Accessibility, Gene Expression and Classification of Tumors

ATAC-Seq datasets were downloaded from TCGA-BRCA project in the National Cancer Institute’s (NCI) Genomic Data Commons (GDC) (18). Datasets were downloaded as bam files, sorted, and read count normalized with DeepTools (v3.1.3) (bamCoverage -bs 10) (19). MACS2 (v2.1.4) was used for peak calls (bdgpeakcall -c 35 -g 100 -l 100) (20). A union peak set was generated containing all peaks across datasets (n=245133), and accessibility in these regions was scored for all tumors. Gene expression datasets were also downloaded from the TCGA-BRCA project. The expression files were downloaded as tables and matched to ATAC-Seq with Case ID. All 1222 expression files available in the TCGA-BRCA project were also combined into a union expression table. Tumor stage and IHC subtype were extracted from the TCGA-BRCA project in the NCI’s GDC. PAM50 subtype (21), histological subtype (22), and general patient demographics (22) data were obtained in cBioPortal (23, 24).

### Unsupervised Dimensional Reduction and Clustering

The union peak table (described above) was uploaded into R and scores were normalized by ranking regions from minimum to maximum accessibility for each tumor. This table was then input into UMAP package (n_neighbors=10) (25). UMAP output three tumor groups by agnostically grouping tumors based on similarities in chromatin accessibility patterns. To identify regions where accessibility differences occurred, log2FC values were calculated from a region’s average accessibility within a tumor group compared with its accessibility in all other tumors. Signatures 1, 2 and 3 consisted of regions with a log2FC greater than 2.5 for groups 1, 2 and 3, respectively. Tumors were considered individually rather than as replicates, and therefore significance measurements were not assessed in defining divergent accessibility or gene expression groups.

### Data Visualization

The pheatmap package in R was used to create heatmaps of chromatin accessibility and gene expression, annotated by tumor characteristics. The ggplot2 package was used to create scatterplots and superimpose characteristics, such as cancer type, on UMAP plots. Integrative Genomics Viewer (IGV) (26) was used to visualize chromatin accessibility in tumor groups and stages. DeepTools was used to create heatmaps of accessibility and ChIP-Seq binding across regions. The Hg38 genome assembly was used.

### Annotation of Chromatin Signatures and Gene Ontology Analyses

HOMER (v4.10) was used to annotate and find motifs enriched in each chromatin signature (see above) (27), and group accessibility trends at those motifs were subsequently determined.

Gene ontologies for chromatin regions were determined with GREAT, which associates regions to any gene whose TSS is within 1000 kb (28). Gene ontologies for genes from divergent gene expression analyses were determined with Enrichr (29, 30).

### Analysis of MCF-7 ChIP-Seq

Encode was used to download ChIP-Seq data from the MCF-7 cell line for FOXA1 (ENCSR126YEB), H3K27ac (ENCSR752UOD), H2A.Z (ENCSR057MWG) and ER (ENCSR463GOT) (31, 32). BigWig files of log2FC over control were downloaded from the ENCODE portal with the following identifiers: ENCFF795BHZ (FOXA1), ENCFF063VLJ (H3K27ac), ENCFF589PLM (H2A.Z), and ENCFF237WTX (ER).

### GSEA

Using the union expression dataset, tables of tumors in the top and bottom decile of *ANP32E* expression were generated. In order to associate gene ontologies with *ANP32E* expression, the average gene expressions of the top and bottom deciles were input into GSEA, which then converted normalized counts data to ranked lists for enrichment scoring (33, 34). To isolate this effect from ANP32E’s association with Basal-L tumors, we sought to eliminate the Basal-L subtype. Using expression of *FOXA1* and *GATA3*, two PAM50 markers, we removed the tumors that were in the bottom quartile of expression for both genes. Testing this method on the 74 known tumors, this results in 14 tumors being eliminated. 10 of the 12 known Basal-L tumors were removed, and 12 of the 14 tumors removed were in group 1. Since this method was shown to be effective in removing the majority of Basal-L tumors from the sample, we applied it to all tumors in the TCGA-BRCA project. This resulted in removing 112 of the 1222 expression files available. We then repeated the GSEA analysis with this subset of tumors.

Statistical analyses were done with R statistical software (v3.6.3), and p-values obtained are from parametric t-tests. Log2 fold-change values were calculated with a pseudo-count of 1.

## Results

### Patterns of Chromatin Accessibility Segregate Breast Tumors into Distinct Subtypes

Chromatin accessibility has been used for defining cell identities, for establishing tissues of origin, and for measuring developmental cell-state transitions (35–38). We therefore sought to identify similarities and differences in chromatin state comparing between breast tumors. Chromatin accessibility maps were previously generated for 74 primary invasive breast carcinomas using ATAC-Seq, as part of TCGA-BRCA project (2, 39). Sequence data (downloaded from the Genome Data Commons-gdc.cancer.gov) were normalized based on total mapped reads, and enriched ‘peaks’ of high accessibility were identified using MACS2 (20) (245133 union peaks). Uniform Manifold Approximation and Projection (UMAP) (25) was then used to group individual tumors based on chromatin accessibility patterns, wherein tumors segregated into three distinct groups (Fig. 1A), with no obvious differences in demographics between groups (Fig. S1A & S1B). Most chromatin differences occurred along UMAP dimension 2, where tumors within group 1 bore the greatest distinction from groups 2 and 3 (Fig. 1A).

**Figure 1:**
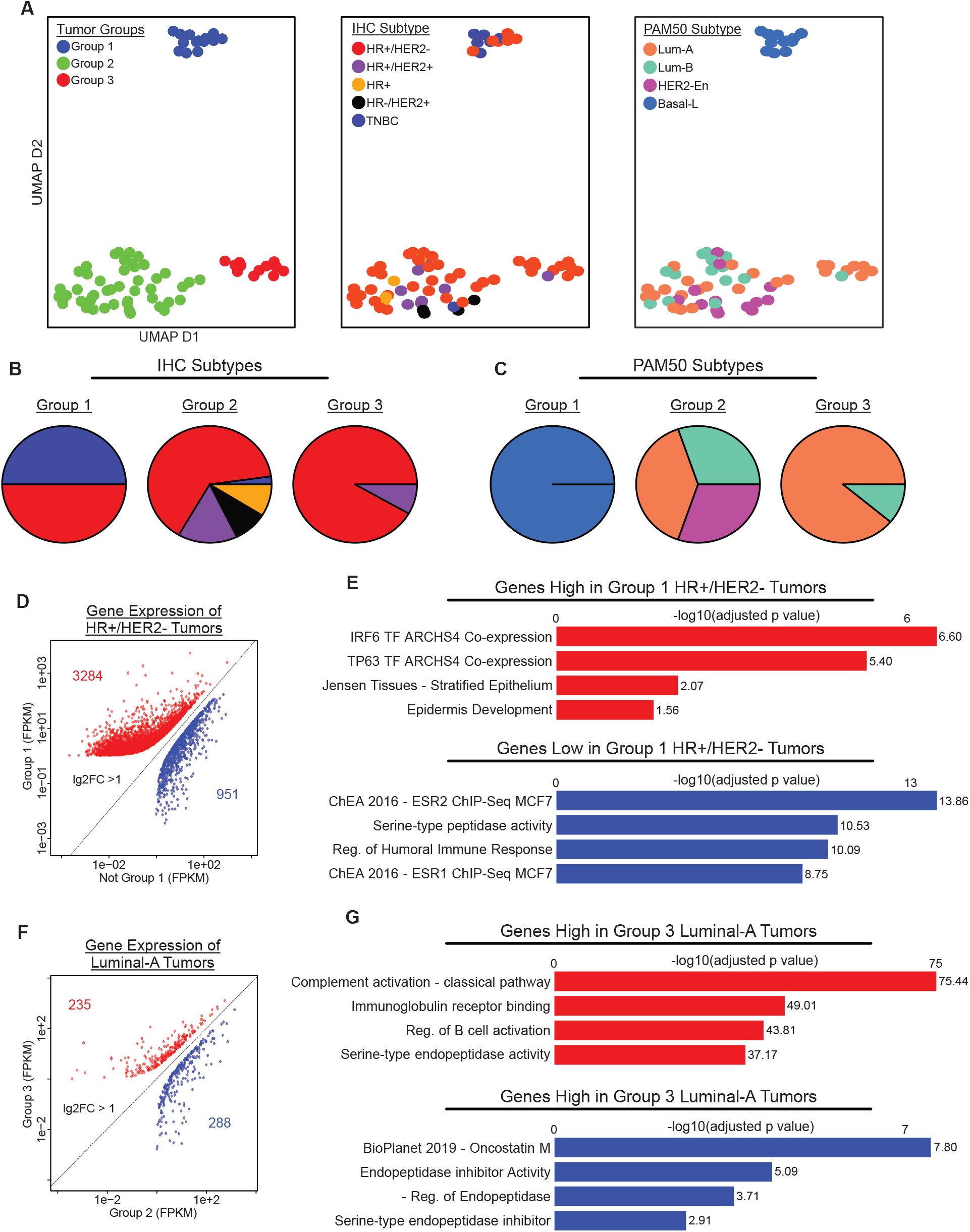
Chromatin Accessibility Distinguishes Breast Cancer Subtypes. A) UMAP dimension reduction plots depicting three distinct groups of tumors, colored by group (n=74), IHC subtype (n=69) and PAM50 subtype (n=65). B-C) Individual pie charts depict groups of tumors based on IHC subtypes (B) and PAM50 subtypes (C), indicating tumor groups distinguish breast cancer subtypes. D-E) Scatterplots depicting genes found to have higher or lower expression in HR+/HER2-tumors in group 1 (n=6) compared to rest (n=40) (D) and in Luminal-A tumors in group 3 (n=8) compared to group 2 (n=17) (E). F-G) Bar charts depicting significance of gene ontology results from Enrichr, investigating genes found to have higher and lower expression in HR+/HER2-patients in group 1 compared to rest (F) and in Luminal-A tumors in group 3 compared to group 2 (G). Adjusted p-values obtained within Enrichr.

We next compared the chromatin-based UMAP classification with existing IHC and PAM50 classifications. We found that group 1 included nearly all TNBC and all Basal-L tumors, whereas all HER2+ tumors (either by IHC or PAM50 classification) were within group 2. HR+/HER2-tumors, however, were distributed across all three groups (Fig. 1A – middle). Perhaps not surprisingly, the chromatin-based UMAP classification better reflected the intrinsic tumor subtypes that are based on gene expression (PAM50 classification) rather than the pathologic classification. For example, Basal-L tumors were found within group 1, and nearly all HER2-enriched and Lum-B tumors were within group 2 (Fig. 1A-right). Interestingly however, a subset of Lum-A tumors (and 1 Lum-B) were classified as a distinct set of tumors within group 3 (Fig. 1A-right) (analyzed subsequently).

We further examined the relationship between the chromatin-based classification and other known features, including the frequency of common mutations and the expression of key genes. As expected (given the enrichment of TNBC/Basal-L tumors in group 1) mutations in *TP53* were over-represented in group 1, whereas mutations in *PIK3CA, GATA3* and *CDH1* were underrepresented (Fig. S1C). Likewise, (given the relationship to PAM50 subtype) expression of *FZD7, SOX9*, and MYC was higher for tumors within group 1, whereas tumors in group 2 and 3 had higher expression of *FOXA1* and *GATA3* (Fig. S2A & S2B).

The above data indicated that features of chromatin accessibility may promote discrete tumor phenotypes, but chromatin patterns are nevertheless distinct from gene expression or histopathology-based phenotypes, suggesting that the chromatin differences may represent unique tumor behavior or underlying biology differences. In this regard, two classes stood out. HR+/HER2-tumors, which were distributed across all three groups, and Lum-A tumors, which were split between groups 2 and 3. To further investigate the nature of these differences, we compared the transcriptomes of those HR+/HER2-tumors that clustered in group 1 with HR+/HER2-tumors outside of group 1. Rather than grouping tumor samples as ‘replicates’, we compared average gene expression levels between UMAP groups and assessed statistical significance in downstream steps. Using this approach, we identified more than 4000 genes with divergent expression (Log2FC >1) (Fig. 1D). As expected, (based on results in Fig 1A) gene ontology (GO) analysis indicated that genes involved in hormone signaling tended to be under-expressed in the HR+/HER2-tumors in group 1 relative to those in the other two groups (Fig. 1E). Among these, expression of *ESR1, PR*, and *ERBB2*, as well as androgen receptor (*AR*) (Fig. S2C) showed reduced expression in group 1 tumors relative to those in groups 2 and 3, and these HR+/HER2-tumors were more similar in the expression of these genes to group 1 tumors classified as TNBC by IHC. These data suggest that chromatin-based classification may be a more accurate reflection of tumor phenotype, and that differences in classification may reflect heterogeneity of HR protein expression, variation in how (low vs. no) HR protein expression is stratified by different sites/pathologists, particularly for ER, and/or differences in mRNA vs. protein-based determination. Consistent with this idea, *ESR1* expression in group 1 HR+/HER2-tumors was not only lower than that in HR+ tumors from other groups, but also exhibited greater variation than tumors determined to be TNBC (i.e. and thus ER-) (Fig. S2C).

As noted above, our chromatin-based classification distinguished a subset of Lum-A tumors (8 of 24) as a distinct group (group 3). There were no apparent differences in the expression of the classic biomarker genes (*ESR1, PR, ERBB2*) or AR (Fig. S2D) between Lum-A tumors in groups 2 vs. those in group 3. We identified 523 genes with divergent expression between group 2 vs. group 3 Lum-A tumors (Fig. 1F). Interestingly, these differences largely reflected dysregulation of genes involves in humoral immune response and inflammatory pathways (which were enriched and depleted) respectively in Lum-A tumors within group 3 (Fig. 1G). Taken together, these data suggest that the chromatin state differences in breast cancer occur largely in Basal-L tumors (as compared with non-Basal-L tumors) and within a distinct subset of Lum-A tumors, potentially resulting from immune evasion (40).

### Accessibility Differences at a Subset of “Signature” Regions Underlie Tumor Groups

To gain further insight into the factors driving group classification, we identified the genomic regions where high levels of accessibility were present for tumors within each respective group, as compared with all other tumors (Log2FC>2.5). This enabled us to define a set of accessible loci (signature regions) which independently partitioned tumors in a manner nearly identical to UMAP grouping (Fig. 2A & 2B). Interestingly, the signature sites for all 3 groups tended to be further away from the nearest annotated TSS (Fig. S3A) and less CpG rich (Fig. S3B), as compared with randomly-selected accessible peak regions, suggesting that they might represent distal regulatory elements. GREAT analysis (28) was then used to annotate the signature regions to all genes whose TSS were within 1000 kb in either direction (Fig. S3). Analysis of the genes near signature 1 sites indicated they were involved in exocrine gland development, which was not surprising given that all tumors within group 1 were Basal-L, and are therefore thought to arise from precursor cells within the basal layer of mammary exocrine glands. By contrast, signature 2 sites were located nearest to hormone responsive genes, consistent with the abundance of HR+ and Lum-A/B tumors in this group. Interestingly, genes associated with signature 3 sites were enriched in functions involved in cell metabolism (Fig. S3E), suggesting that a unique metabolic program may distinguish tumors in this group from those that otherwise bear a Lum-A gene expression signature.

**Figure 2:**
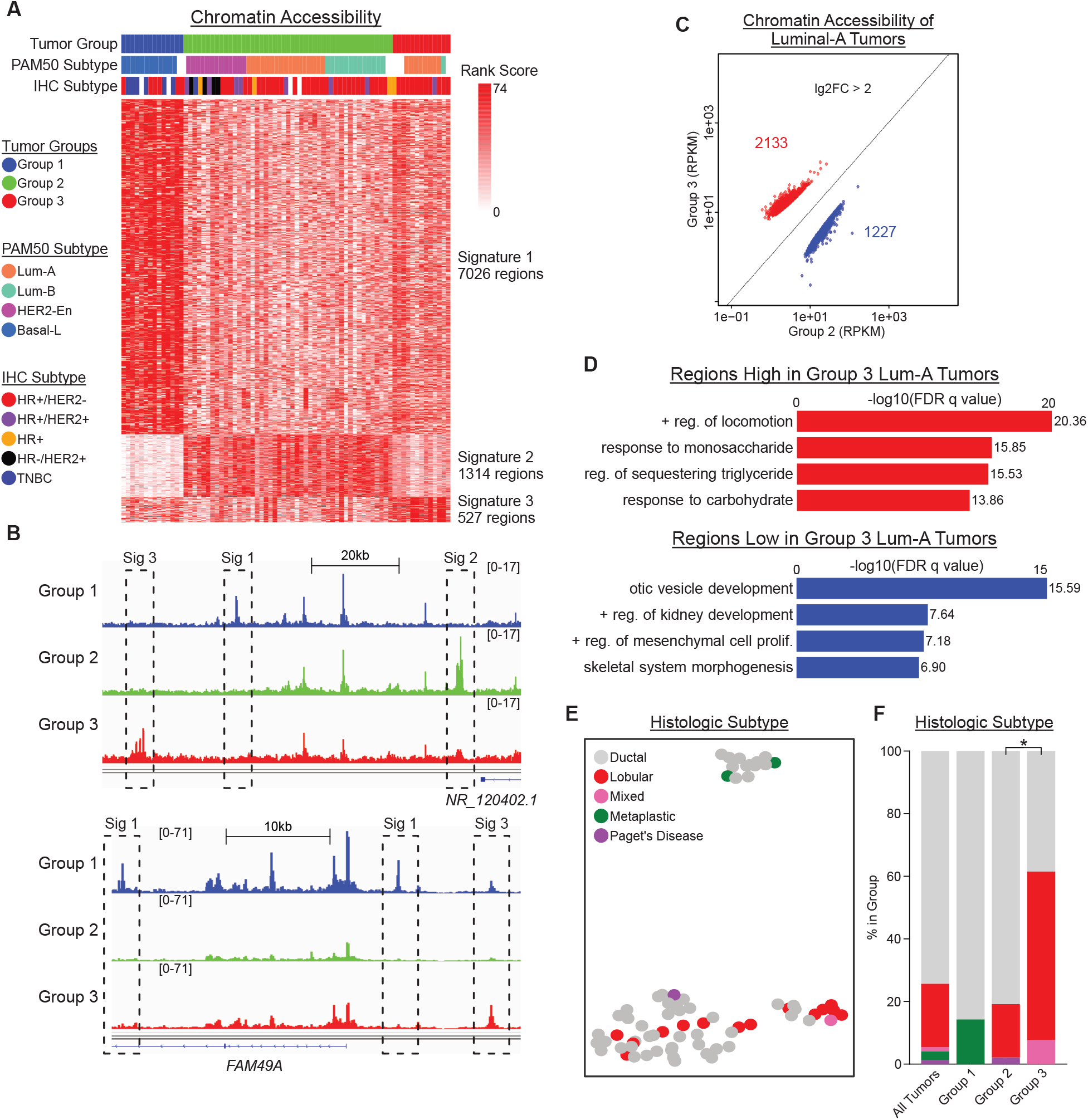
Distinct Chromatin Accessibility Signatures Associate with each Tumor Group. A) Heatmap showing 3 groups of chromatin regions, each showing greater accessibility in their respective tumor group compared to the rest (Lg2FC > 2.5). B) Screenshots from IGV depicting average accessibility of tumor groups in regions within each chromatin signature. C) Scatterplot displaying regions found to have higher or lower accessibility in Luminal-A tumors in group 3 (n=8) compared to group 2 (n=17). D) Bar charts depicting significance of gene ontology results from GREAT, investigating genes nearby (<1000 kb) regions found to have higher and lower accessibility in group 3 Luminal-A tumors compared to group 2. FDR q-values obtained within GREAT. E-F) UMAP plot (E) and stacked barplot (F) showing the distribution of cancer types between each tumor group. P-value in F obtained from Chi-squared test within cBioPortal.

As noted, the chromatin profiles segregated Lum-A tumors into two groups (Lum-A, group 2 vs group 3) (Fig. 1C). To gain further insight into the potential factors segregating these tumors, we applied a similar approach as above to identify regions of chromatin accessibility that differed between Lum-A tumors in group 2 and group 3, including sites outside our defined signature regions (log2FC>2, n=3360) (Fig. 2C). Interestingly, GREAT gene ontology analysis (genes within 1000 kb) revealed that regions of greater accessibility in group 2 Lum-A tumors (lower in group 3) were annotated to genes involved in development and morphogenesis, whereas regions with greater accessibility in group 3 were annotated to genes involved in carbohydrate metabolism (Fig. 2D). These results provide further indication that a subset of Lum-A tumors may be programmed in a metabolically distinct manner based on differences in chromatin state. Both the subset of Lum-A tumors in group 3, and group 3 tumors in general, were distinguished by features associated with immune (Fig. 1G) and metabolic regulation (Fig. 2D), similar to gene expression characteristics previously identified as distinguishing subsets of invasive lobular carcinomas (ILC) which exhibit a Lum-A intrinsic gene expression pattern upon PAM50 subtyping (41–43). To investigate this further, we overlayed the tumor histology information extracted from the TCGA metadata for each tumor in the dataset along with the UMAP classification (Fig. 2E). We found that indeed, ILC was over-represented in group 3, relative to groups 1 and 2 (Fig. 2F). ILC is also characterized by high rates of CDH1 mutation, which were also found to be somewhat overrepresented in group 3 tumors (Fig. S1C). However, many (8 of 14) lobular carcinomas also distributed to group 2, indicating that chromatin state differences occurred in a subset of lobular carcinomas, many of which were classified previously as Lum-A (Figs 1A & 1G).

### Accessibility at FOX and SOX binding sites defines tumor groups

To better understand how chromatin changes might contribute to biologically distinct tumor properties, we next investigated the genomic context of the established signature regions. The gene-distal nature of these signature regions (Fig. S3A) suggests that they might function as intergenic regulatory sites. To investigate this possibility, we used HOMER (27) to identify DNA sequence motifs enriched in the signature regions, representing potential transcription factor binding sites. Several motifs were found to be enriched (Supplemental Table 1), as compared with background regions (consisting of 5000 randomly selected, similarly sized genome-wide accessible sites). SOX factor binding motifs were most enriched in signature 1 regions, FOX factor motifs were the most enriched in signature 2 regions, and CEBP motifs were the most enriched in signature 3 regions (Fig. 3A). To confirm that accessibility differences occurred directly over these candidate motifs, we next mapped motif locations within signatures 1, 2, and 3, and assessed accessibility levels at these sites (Fig. 3B – top). Indeed, tumors in group 1 had, on average, greater accessibility at SOX motifs, group 2 tumors had the highest accessibility at FOX motifs, and group 3 tumors had the highest accessibility at CEBP motifs.

**Figure 3:**
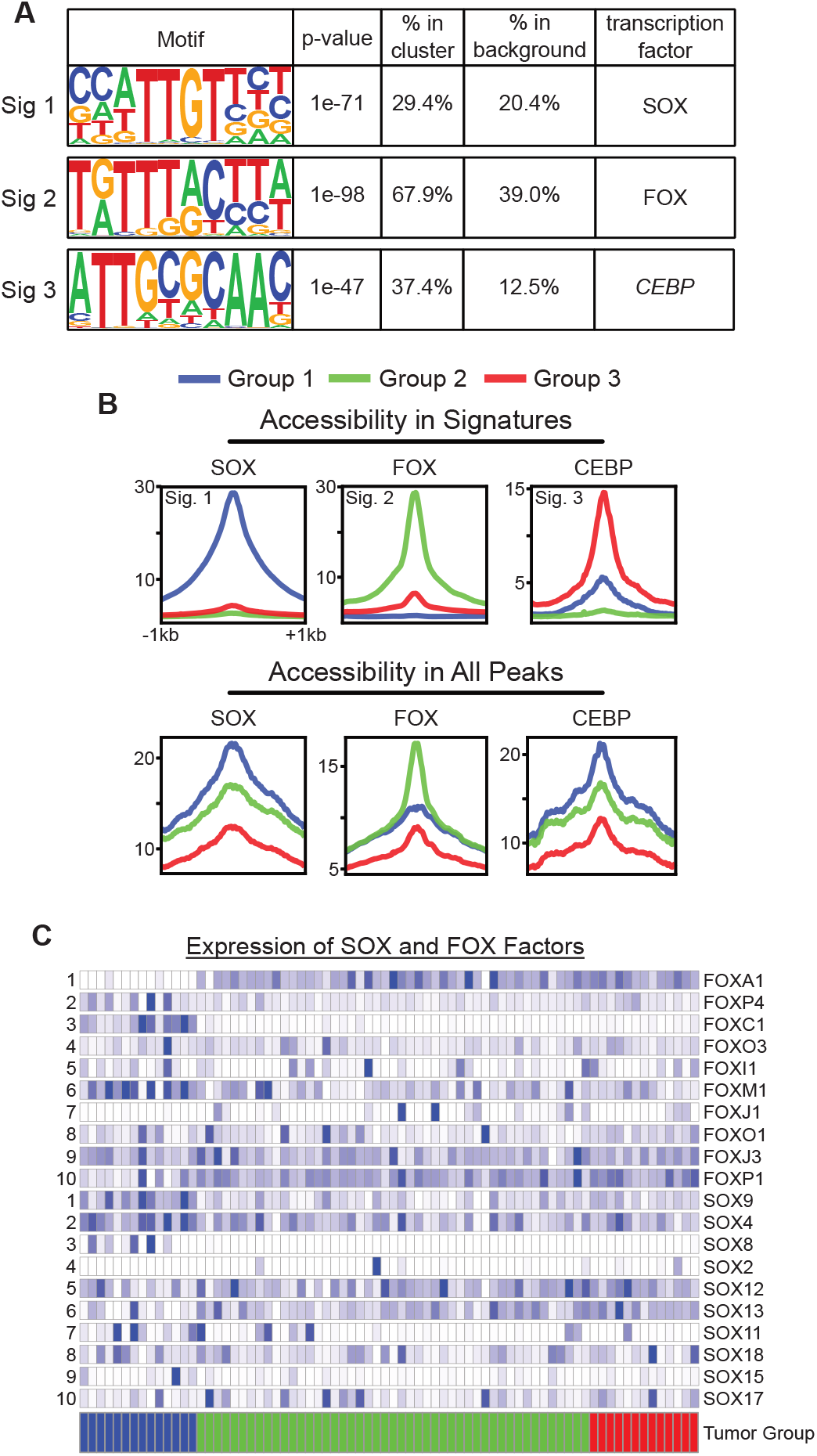
Accessibility at FOX and SOX Binding Sites Define Tumor Groups. A) Table displaying top motif result from HOMER for each chromatin signature. Due to similarity across SOX and FOX binding motifs, we refer to SOX6 simply as SOX, and FOXM1 simply as FOX. P-values obtained within HOMER. B) Profile plots depicting average accessibility of tumor groups in motif regions, both within the motif’s respective signature (top) and in all accessible peak regions in tumors (bottom), indicating that group 1 and 2 tumors show increased accessibility at SOX and FOX motifs, respectively. We again use the SOX6 motif to represent SOX motifs, and the FOXM1 motif to represent FOX motifs. C) Heatmap showing expression of SOX and FOX factors across tumor groups.

Having identified DNA motifs enriched within each signature region, we next asked whether these motif locations were more broadly accessible throughout the genome, even when motifs were located outside our signature regions (Fig. 3B – bottom). Indeed, we found that genome-wide, accessibility at SOX motifs was highest in group 1 tumor samples, and at FOX motifs, accessibility was highest in group 2 samples (hormone positive tumors). Interestingly, the accessibility at SOX, FOX, and CEBP sites was lower for group 3 tumors, when compared with groups 1 and 2, suggesting that additional factors may underlie accessibility of the group 3 tumors. We next compared the levels of gene expression to determine which among the FOX and SOX family transcription factors might be involved. Here we found that group 1 tumors tended to express high levels of *SOX9, FOXC1*, and *FOXM1* relative to tumors in groups 2 and 3, whereas group 2 tumors expressed high levels of *FOXA1* (Fig. 3C & S4A). These results are well aligned with published studies indicating that *SOX9* and *FOXC1* play central roles in TNBC (within group 1), whereas FOXA1 status and levels are mediators of outcome and programming among ER+ breast tumors (mostly in group 2) (11, 12, 14).

HR+ breast cancers in groups 2 and 3 expressed higher levels of *FOXA1* (Fig. S4A) and exhibited greater chromatin accessibility over FOX factor sequence motifs (Fig. 3B). Prior studies indicate that FOXA1 function in conjunction with ER to influence enhancer activity and promote pro-metastatic transcriptional programming in breast cancer cell lines (14, 15). We therefore determined the relationship between chromatin grouping and tumor progression. Indeed, we found that average chromatin accessibility levels at the signature regions defining groups 1-3 were associated with tumor stage (Fig. 4A-C). Notably, at signature 2 regions, which are enriched for FOX motifs (Fig. 3A), there was a strong relationship between accessibility and tumor stage, with progressively greater accessibility associated with increasing severity of disease (Fig. 4C & S4B). There was a nonsignificant trend towards decreased accessibility at signature 3 sites with increasing stage of disease. Signature 1 sites showed the greatest accessibility in early stage (stage I and II), suggesting that among Basal-L/TNBC breast cancers such sites may underlie the early stages of disease.

**Figure 4:**
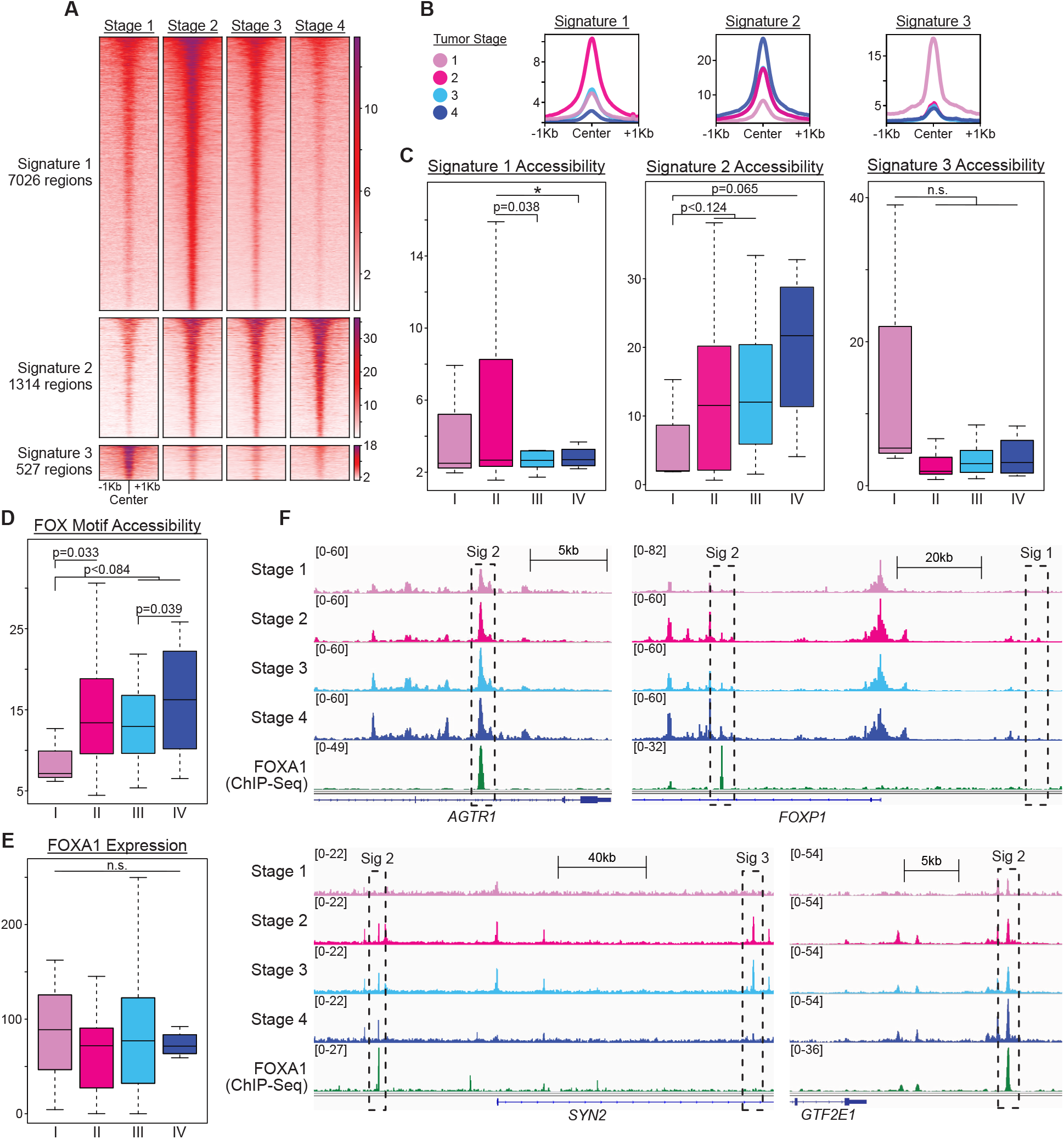
Chromatin Accessibility in FOX motifs and Signature 2 Regions Associate with Tumor Progression Stages. A-B) Heatmaps (A) and profile plots (B) showing accessibility in signatures 1, 2 and 3 across tumor stages. Heatmaps have regions ordered from greatest to least average accessibility across tumor stages. C) Boxplots of accessibility in signatures 1, 2 and 3 across tumor stages, indicating that only signature 2 shows an accessibility trend across stages. D-E) Boxplots comparing accessibility of FOX motifs in accessible peak regions (n=96280) (D) and FOXA1 expression levels (E) across tumor stages. F) Screenshots from IGV depicting average accessibility of tumor stages and FOXA1 binding in MCF-7 cells from ChIP-Seq in regions within each chromatin signature. ChIP-Seq data is Log2FC over control. P-values in C-E obtained from one-tailed parametric t-tests. * is p<0.01, ** is p<0.001, *** is p<0.0001.

The observation that signature 2 regions were positively associated with increasing tumor stage among group 2 tumors (Fig. 4A) was recapitulated in our analysis of accessibility at FOX motifs across all accessible sites for the 74 tumors in the dataset (n=96,280) (Fig. 4D & S4C). Here we found again that accessibility of FOX motifs among all accessible sites is positively correlated with tumor stage, despite no apparent differences in *FOXA1* gene expression between tumors of different stages (Fig. 4E). Similar to what was observed for differences in accessibility at FOX motifs (Fig. 3D), we noted a trend towards greater accessibility at FOX motifs among Lum-A or lobular carcinomas in group 2 vs. group 3. Regardless of whether we focused on Lum-A or lobular tumors, those in group 2 had higher accessibility levels than tumors in group 3 (Fig. S4D) – despite no apparent differences in *FOXA1* expression levels (Fig. S4E).

Cognizant of the previously described relationship between ER and FOXA1 binding at distal enhancer elements in breast cancer cells (as discussed above (14, 15)), we next investigated histone modifications at the accessible signature regions. Using publicly available chromatin immunoprecipitation data from MCF-7 cells (31, 32), we found that FOXA1, ER, and H3K27ac (a marker of active enhancers) are significantly enriched at signature 2 regions, as compared with accessible regions that define groups 1 or 3 (Fig. 5A). This high level of co-occurrence suggests that the co-binding of FOXA1 and ER may underlie the chromatin-based segregation in HR+/Lum-A/lobular carcinomas between groups 2 vs. 3. When considered together, these results suggest that chromatin accessibility at FOX factor binding motifs increases with tumor progression, and this mechanism underlie distinguishing chromatin patterns in group 2 verses group 3 tumors.

**Figure 5:**
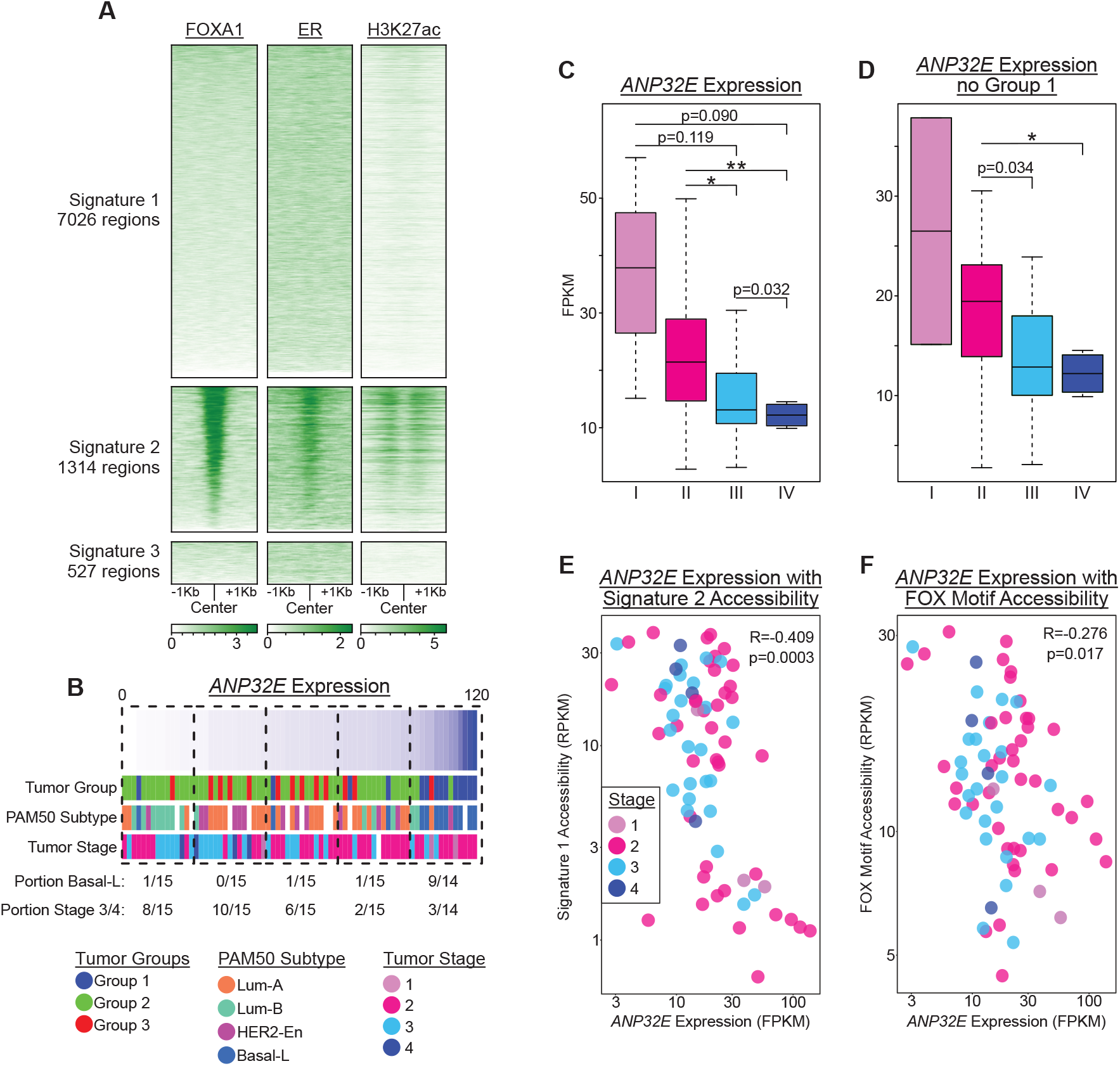
*ANP32E* Expression Levels Associate with FOX Motif Accessibility and Tumor Stage. A) Heatmaps showing binding of FOXA1, ER, and H3K27ac in MCF-7 cells within regions from signatures 1, 2 and 3. Data from ChIP-Seq of MCF-7 cells; regions sorted from greatest to least FOXA1 enrichment. B) Heatmap with tumors ordered by *ANP32E* expression and annotated by tumor group, PAM50 subtype and tumor stage, indicating that late-stage tumors group with lower *ANP32E* expression. C-D) Boxplots comparing *ANP32E* expression by tumor stage, both in all tumors with available stage data (n=73) (C) and in only tumors from groups 2 and 3 (n=59) (D), indicating that late-stage tumors have significantly lower expression of *ANP32E*. P-values obtained from one-tailed parametric t-tests. * is p<0.01, ** is p<0.001, *** is p<0.0001. E-F) Scatterplots showing correlation of *ANP32E* expression with tumor’s accessibility in signature 2 regions (E) and with tumor’s average accessibility in FOX motifs across all accessible peak regions (n=96280) (F), with tumors colored by stage. R denotes Pearson correlation coefficient; p-values from Pearson’s product moment correlation coefficient.

### *ANP32E* Expression Levels are Associated with Accessibility at FOX Motifs

The above data indicate that accessibility of FOX motifs is generally associated with tumor stage, but we find little evidence for differences in *FOXA1* (or *ESR1*) expression levels between tumors of different stages, suggesting that additional factors may contribute to accessibility at FOX binding motifs. Prior studies of HR+ breast cancer cells have demonstrated that the function of FOXA1 is impacted by the local enrichment of the histone variant H2A.Z (44, 45). H2A.Z accumulates at estrogen response elements that are bound by FOXA1 and loss of H2A.Z impairs both FOXA1 binding and polymerase recruitment. ANP32E is a chromatin chaperone that regulates the genomic localization of H2A.Z to control locus-specific chromatin state dynamics (16). In recent work, we showed that ANP32E antagonizes H2A.Z installation, such that ANP32E loss causes a global increased H2A.Z enrichment, heightened chromatin accessibility and amplified TF-binding at open sites, in cultured mouse fibroblasts (16). ANP32E may function similarly in breast tumors, influencing the binding of key oncogenic transcription factors, like FOXA1. Therefore, we investigated the relationship between *ANP32E* expression, chromatin accessibility, and tumor characteristics across the chromatin-defined tumor groups (Fig. 5B). We found that *ANP32E* expression was generally higher in group 1 tumors than groups 2 and 3 (Fig. 5B & S5A), and consistent with a prior report looking at protein levels (17), ANP32E was significantly higher in Basal-L tumors (within group 1) than the other PAM50 subtypes (Fig. 5B & S5B). Moreover, the levels of *ANP32E* expression tended to stratify tumors by stage, wherein early-stage (I, II) tumors had the highest levels of *ANP32E* expression and late-stage (III, IV) tumors had the lowest levels (Fig. 5B, 5C & S5C). This association was maintained even when group 1 tumors were excluded (Fig. 5D & S5D), indicating that *ANP32E* levels may be functionally involved in cancer progression independent of tumor subtype.

We next evaluated the relationship between ANP32E expression and accessibility. We found that accessibility at signature 2 regions (Fig. 5E) or all accessible FOXA1 motifs (Fig. 5F) were significantly anticorrelated with levels of *ANP32E* expression across all tumors (signature 2: R= -0.409, p=0.0003; FOX motifs: R= -0.276, p=0.017), suggesting that ANP32E may indeed function as a negative regulator of chromatin accessibility at these sites. Notably, signature 2 regions had the highest levels of H2A.Z in MCF-7 cells (Fig. S5E), and the negative relationship between *ANP32E* expression and accessibility was specific to signature 2 (Fig. S5F & S5G).

### ANP32E Expression Levels are Associated with DNA Replication and Immune Response

Based on our findings that reduced *ANP32E* expression levels associated with tumor stage progression, we next sought to determine the relationship between *ANP32E* expression and tumor phenotype, using the tumor transcriptome as a read-out. For this analysis, we first identified tumors in the top and bottom deciles of *ANP32E* expression (n=123) among all breast tumors from the TCGA-BRCA project for which RNA-Seq data were available (n=1222). Then, we used GSEA to investigate the ontologies of these transcriptomic shifts associated with *ANP32E* expression levels. We found that high *ANP32E* expression was associated with increased expression of genes involved in the immune response (Fig. 6A) and to a lesser extent DNA replication (Fig. S6A). Consistent with this idea, *KI67* expression, a marker of cellular proliferation (46), was highest in group 1 tumors (representing all Basal-L and most TNBC tumors) (Fig. 6B), and *ANP32E* and *KI67* levels were positively correlated across all samples analyzed, but not after removing group 1 (Basal-L) tumors (Fig. 6C). Conversely, low *ANP32E* expression was associated with increased expression of genes involved in separate developmental processes (eg. ‘Keratinocyte Differentiation’ and ‘Cilium Movement’). To test whether the observed gene expression associations were driven by differences between Basal-L and non-Basal-L tumors, we repeated these analyses exclusively assaying tumors classified as non-Basal-L (see methods). Here again we found that high *ANP32E* expression was associated with genes involved in DNA replication and immune response (Fig. 6D & S6B), indicating that the observed GSEA results were not simply the outcome of tumor subtype gene expression differences.

**Figure 6:**
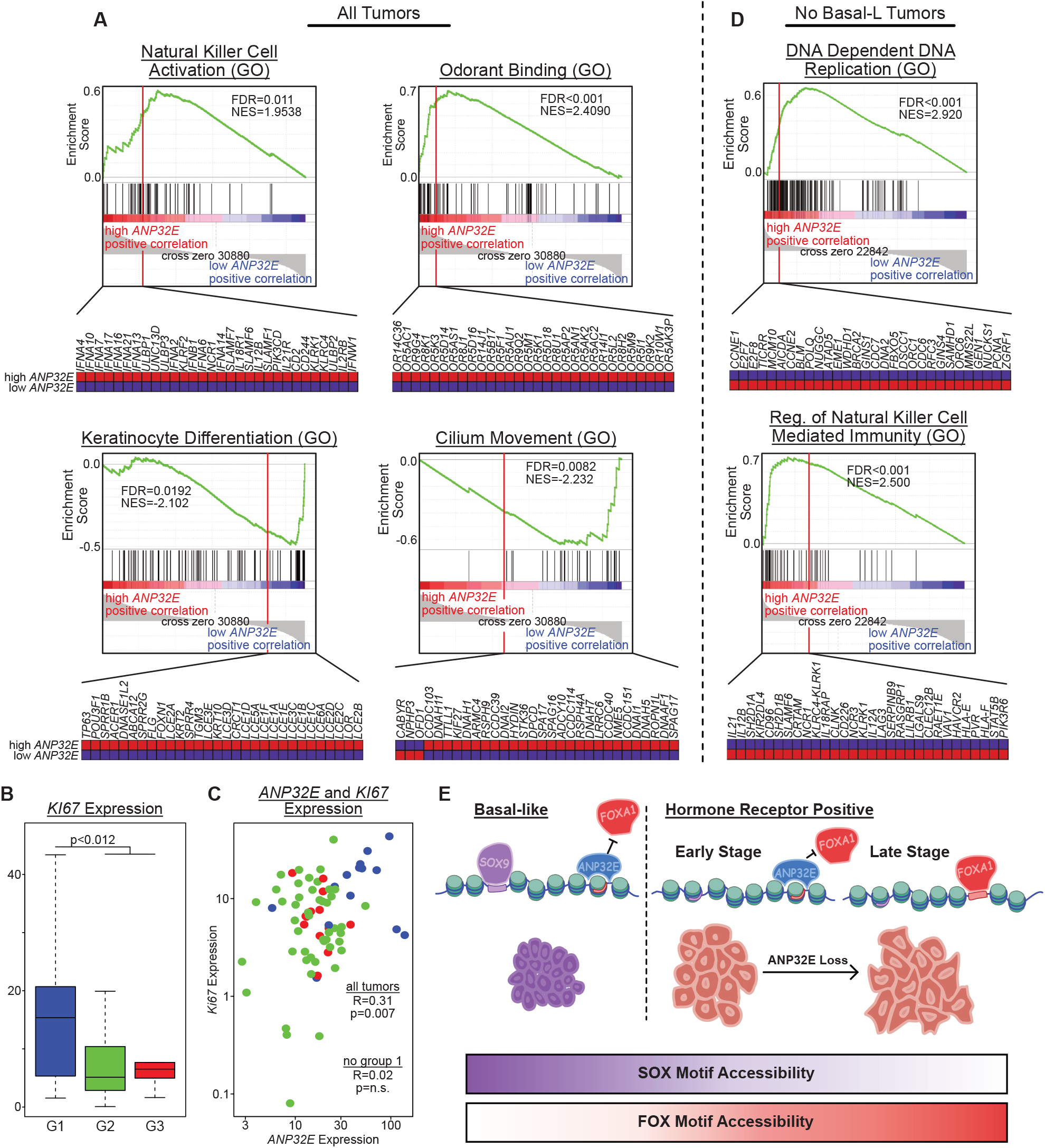
*ANP32E* Expression Levels Associate with Distinct Expression Profiles. A) GSEA plots depicting gene ontology associations with high and low *ANP32E* expression levels for all tumors with RNA-seq data in the TCGA-BRCA project (n=1222). FDR values and normalized enrichment scores (NES) obtained within GSEA. B) Boxplot of *KI67* expression across tumor groups. P-value from one-tailed parametric t-test. C) Scatterplot showing the association between *ANP32E* and *KI67* expression levels, with tumors colored by tumor group. R denotes Pearson correlation coefficient; p-values from Pearson’s product moment correlation coefficient (done for all tumors, and all tumors excluding group 1). D) GSEA plots for all tumors excluding Basal-L (n=1110). E) Model displaying the association of *ANP32E* expression levels with multiple characteristics of breast cancer.

Taken together, these results suggest that ANP32E may generally function to restrict chromatin changes at the beginning stages of tumor development, and loss of ANP32E promotes tumor progression by enabling more aggressive cancer. In this regard ANP32E may act to ‘lock in’ a defined chromatin state, and when tumor cells transition to later stages of cancer progression, ANP32E becomes downregulated, leading to increased chromatin accessibility at a defined set of gene regulatory regions, including sites where H3K27ac and H2A.Z are enriched, enhancer elements, and FOXA1 binding sites.

## Discussion

We set out to investigate how differences in chromatin state across separate breast tumors coincided with unique characteristics of cancer biology, and to investigate whether differences in chromatin patterns could provide insight into new cancer mechanisms. To test whether chromatin accessibility patterns differed in a biologically meaningful manner, we took an unsupervised approach, using a dimensional reduction method (UMAP) to group tumors based only on chromatin differences. With this approach, 74 breast cancer tumors were grouped into three distinct UMAP categories. Supporting the validity of our UMAP approach, we found that differences in chromatin patterns associated with several known breast cancer features, including IHC marker status (Fig. 1B), PAM50 subtype classification (Fig. 1C), and histological classification (Fig. 2F). We also uncovered several novel chromatin associations. For example, our UMAP analysis indicated that 6 HR+/HER2-tumors were more similar to TNBC tumors (Fig. 1A), and these tumors were distinct from other HR+/HER2-tumors. Further characterization revealed that these 6 samples, along with TNBC samples, were classified as Basal-L, suggesting that the chromatin state of Basal-L tumors drove the UMAP segregation patterns, rather than chromatin associations with IHC marker status. Differences in tumor heterogeneity may contribute to these differences in grouping and IHC status. For example, HR+/HER2-tumors with non-uniform IHC staining may be more similar to TNBC tumors when considered in aggregate than homogeneously stained HR+/HER2-tumors. Another interesting possibility is that a subset of HR+/HER2-tumors may be mechanistically more similar to TNBC-like tumors, perhaps explaining why a subset of HR+/HER2-tumors are resistant to hormone therapies (47). More comprehensive and longitudinal studies of breast cancer, measuring chromatin state changes along with IHC status and gene expression profiling, will help in establishing which of these possibilities account for the observed UMAP grouping. In the future, additional diagnostic tests of HR+/HER2-tumors may be necessary to assess intrinsic cell-type of origin, potentially strengthening predictions of therapy response.

We also found chromatin differences occurred in a subset of Lum-A tumors, which appeared to have chromatin patterns more similar to non-Lum-A tumors within UMAP group 2 (including Lum-B and HER2-enriched tumors). This subset had reduced expression for genes involved in immune response (Fig. 1G) and reduced accessibility at regions proximal to metabolism genes (Fig. 2D), despite no measurable difference in expression for typical breast cancer markers, such as *PR, ESR1*, and *ERBB2* (Fig. S2E). We observed similar patterns for lobular tumors, which also segregated into two classes (Fig. 2E s& 2F), with a subset of lobular tumors grouping with ductal tumors (within UMAP group 2). Previous studies examining differences in Lum-A carcinomas found that pathways similar to those active in group 3 tumors were also active in ILC (as compared with ductal carcinoma), including immune-related and metabolic pathways (43). In this context, our results suggest that group 3 may represent invasive carcinomas, similar to those described previously (41–43). Indeed, prior studies found that ILC had decreased FOXA1 activity (based on measurements of gene expression and mutation frequency) (41), and in our study, we found lower chromatin accessibility levels at FOXA1 binding sites in group 3 tumors, which we presume to be of this same ILC subtype (Fig. 3B). In sum, our results support a model where loss of FOXA1 activity (and subsequent loss of DNA binding) in luminal tumors distinguishes ILC from other HR+ tumors (presumably occurring within UMAP group 2).

We found the tumors within UMAP group 2 to be particularly interesting, as several distinct cancer subtypes grouped together, indicating that they had quite similar chromatin accessibility patterns despite differences in clinical classifications. Interestingly, FOX motifs were enriched within the genomic loci where accessibility differences distinguished group 2 tumors (Fig. 3A & 3B) and these loci were located distal from gene promoters (Fig. S3A). In MCF-7 breast cancer cells, these regions are bound by FOXA1 and ER, and enriched for H2A.Z and H3K27ac (Fig. S5E & 5A), suggesting that they may function as enhancer elements in HR+ breast cancer tumors. Prior studies have demonstrated that H2A.Z levels at ER binding sites facilitates enhancer activation and FOXA1 binding in this type of (HR+) breast cancer cell (44, 45). We and others previously demonstrated that H2A.Z is a negative regulator of DNA methylation (48– 50), and accordingly, lower DNA methylation levels are known to occur at enhancers bound by FOXA1 and ER in luminal tumors (compared with basal tumors) (51). Additionally, increased FOXA1 activity has been shown to function in the activation of pro-metastatic cellular programming (14). Taken together, these results suggest that increased H2A.Z levels at enhancers in luminal tumors may promoter increased accessibility, improved FOXA1 binding, and amplified enhancer activity, potentially driving tumors toward a more metastatic cellular program without changing FOXA1 expression levels (Fig. 6E).

The histone chaperone ANP32E has previously been shown to control H2A.Z levels at thousands of vertebrate gene regulatory regions, including enhancers (16, 48, 52, 53). We previously found that ANP32E functions in mouse cells to control genome-wide chromatin accessibility through regulation of H2A.Z patterns (16). Based on this mechanism, differences in ANP32E levels among breast tumors may lead to differences in H2A.Z enrichment, causing chromatin accessibility differences, and ultimately impacting transcription factor binding events. In the context of this study, we do indeed find that ANP32E expression levels differ among tumors, and these differences are anticorrelated with chromatin accessibility at FOX factor binding sites. Interestingly, accessibility at these same sites tends to increase in later-stage tumors (stage III, IV), compared with earlier stages (stage I, II), suggesting that selective opening of signature 2 regions, and FOX binding in particular may function to promote tumor progression. In this regard, ANP32E levels in HR+ tumors may specifically restrict chromatin accessibility at FOX factor motifs (Fig. 6E). Additional mechanistic studies of ANP32E, H2A.Z and its role in FOX factor binding in the context of HR+ breast cancer will be necessary to further investigate this possibility.

It is important to note that our study investigated accessibility data from primary tumor samples only. In this context, our ability to identify significant correlations between stage at resection and chromatin accessibility suggests that changes in the chromatin state of the primary tumor may precede, and/or be predictive of, the propensity for tumor progression and/or metastatic spread. We therefore propose a model in which ANP32E has two separate functions in breast cancer, depending on tumor subtype or the differentiation state of the cell of origin. In Basal-L/TNBC tumors, largely believed to arise from a more stem-like multipotent progenitor, high levels of ANP32E ‘lock-in’ a pattern of accessible chromatin that favors proliferation and self-renewal, while in HR+ breast tumors, arising in a more differentiated luminal progenitor, ANP32E supports the maintenance of luminal identity and hormone responsiveness by restricting FOXA1 binding at estrogen response elements (Fig. 6E). In this latter setting, the loss of ANP32E expression may lead to increased FOXA1 binding, relaxation of cellular programming, and progression to a hormone-resistant state. Indeed, factors affecting the balance of ER and FOXA1 binding to estrogen response elements, such as forced overexpression of FOXA1, may promote expression of genes involved in metastasis and endocrine-resistant breast cancers (14). Future studies addressing the role of ANP32E, H2A.Z, and their role in FOX factor binding in the context of HR+ breast cancer are necessary to further investigate this possibility.

## Acknowledgements

The results published here are in whole or part based upon data generated by the TCGA Research Network: https://www.cancer.gov/tcga. TCGA-BRCA dbGaP Study Accession: phs000178.v11.p8.

