## Supplemental Figures for "Chromatin Patterns Distinguish Breast Tumor Subtypes and Disease Progression in Association with *ANP32E* levels"

Supp. Fig. 1

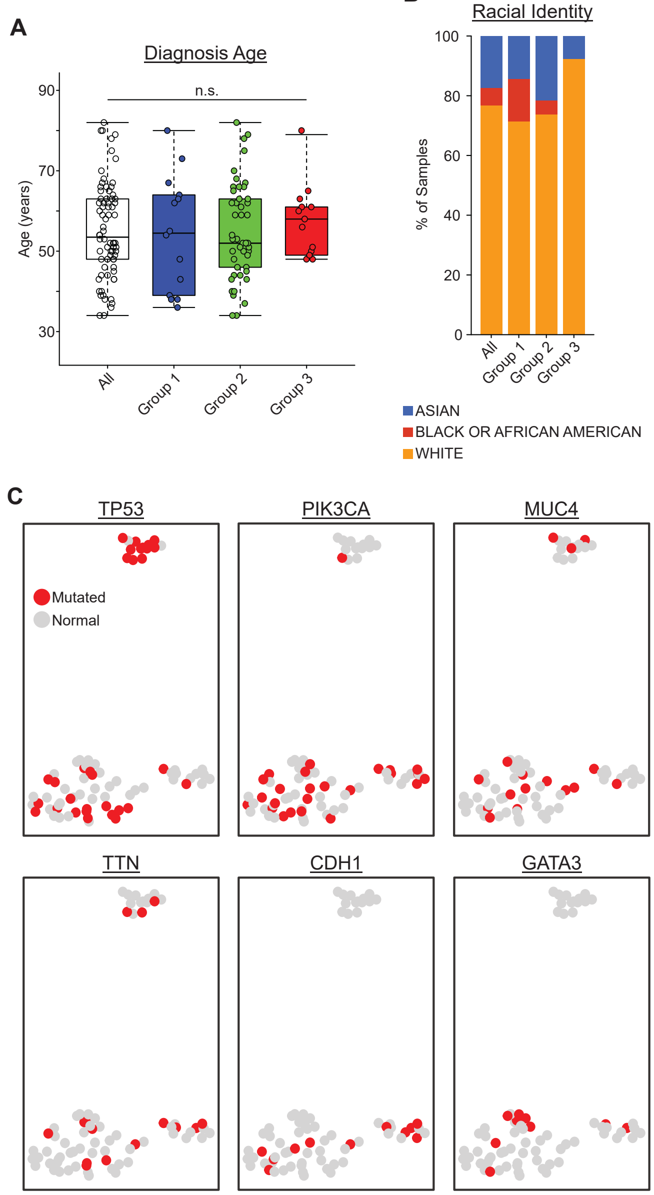


Figure 1 supplement:

A-B) For patients whom tumors were obtained from, boxplot showing age at diagnosis (A) and stacked barplot displaying racial identity (B). Significance values obtained within cBioPortal with Kruskal-Wallis test. C) UMAP plots colored by tumor's mutation status for commonly mutated genes in the TCGA-BRCA project.

**Supp. Fig. 2**

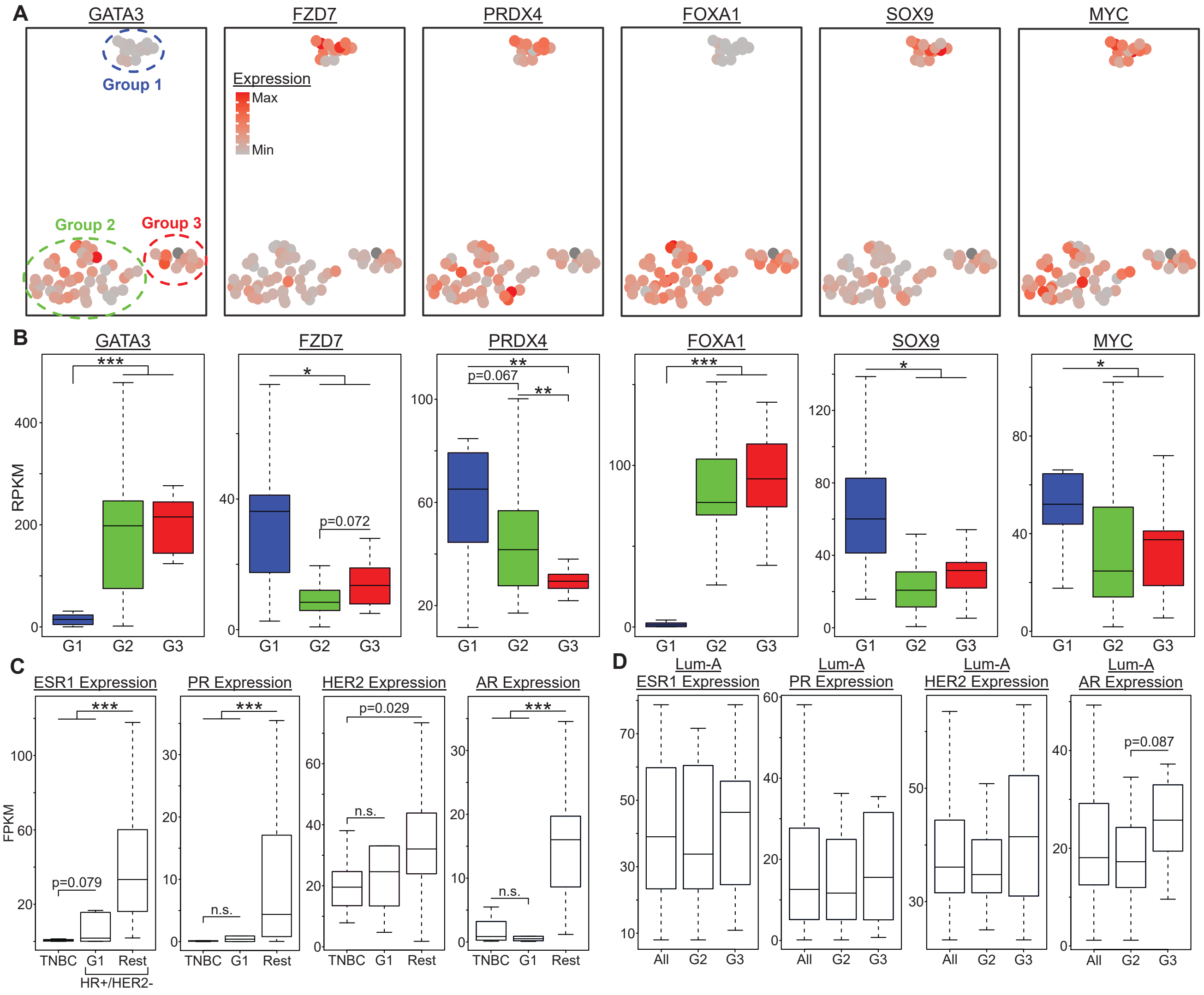


Figure 2 supplement:

A-B) UMAP dimension reduction plots colored by tumor's expression for PAM50 genes (A) and boxplots comparing those gene expressions by tumor group (B). C-D) Boxplots comparing gene expression of hormone receptors in TNBC tumors (n=7) and HR+/ HER2- tumors separated into group 1 (n=6) and rest (n=40) (C) and in Luminal-A tumors separated into all (n=25), group 2 (n=17) and group 3 (n=8) (D). P-values in B-D obtained from one-tailed parametric t-tests. * is p<0.01, ** is p<0.001, *** is p<0.0001

**Supp. Fig. 3**

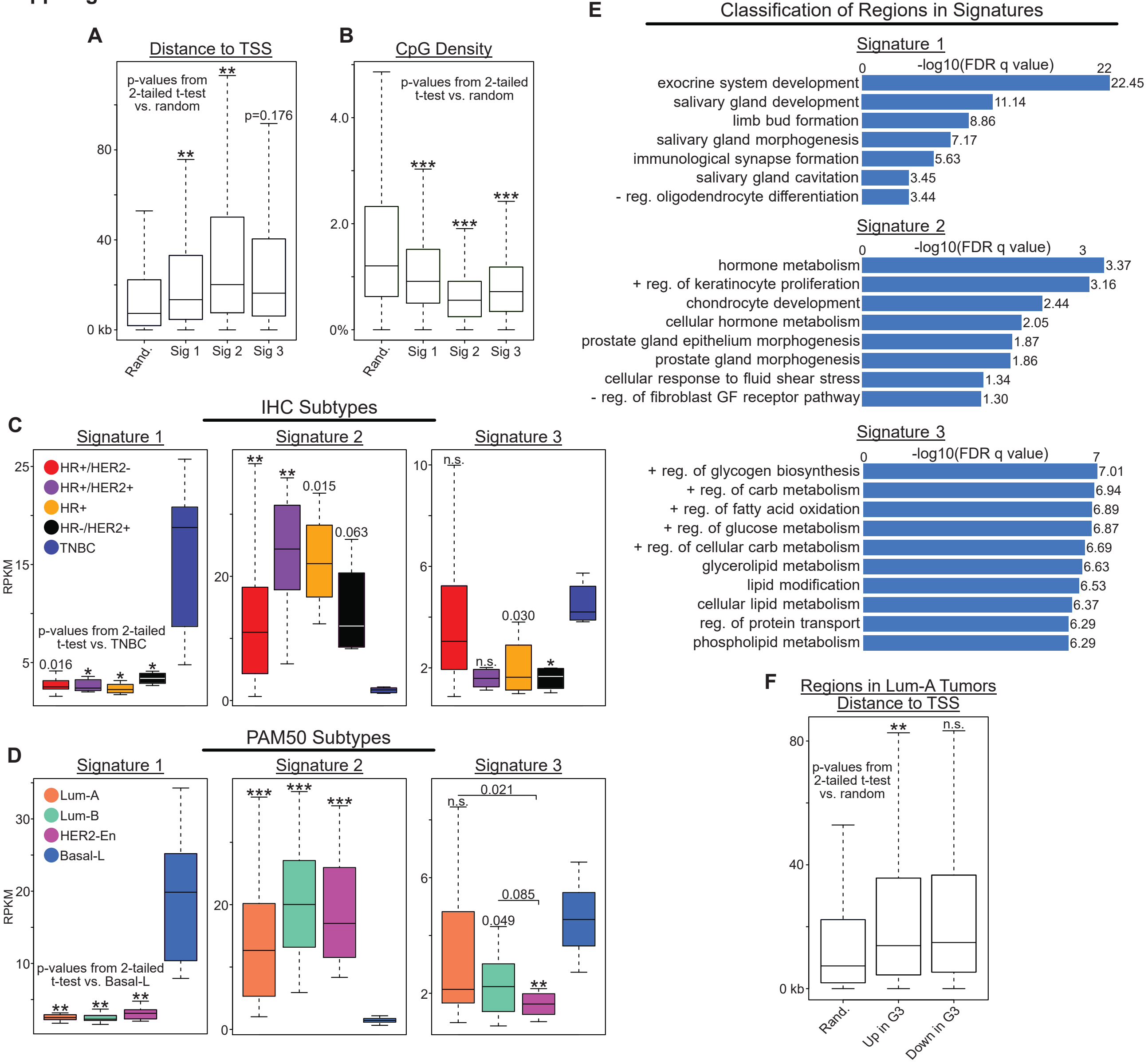


### Figure 3 Supplement:

A-B) Boxplots comparing chromatin signatures by regions' distance to transcription start sites (A) and CpG density (B), with random accessible regions from the genome as a control. P-values obtained from two-tailed parametric t-tests. C-D) Boxplots comparing chromatin accessibility of tumors by IHC (C) and PAM50 (D) subtypes, for regions in signatures 1-3. P-values not noted with brackets obtained from two-tailed parametric t-tests, while one-tailed parametric t-tests are noted by one-tailed brackets. E) Bar charts depicting significance of gene ontology results from GREAT, investigating nearby (<1000 kb) genes to regions within each chromatin signature. . FDR q-values obtained within GREAT. F) Boxplot of regions' distance to transcription start sites for regions found to have higher and lower accessibility in group 3 Luminal-A tumors (from Fig 2C), with random accessible regions from the genome as a control. * is p<0.01, ** is p<0.001, *** is p<0.0001.

**Supp. Fig. 4**

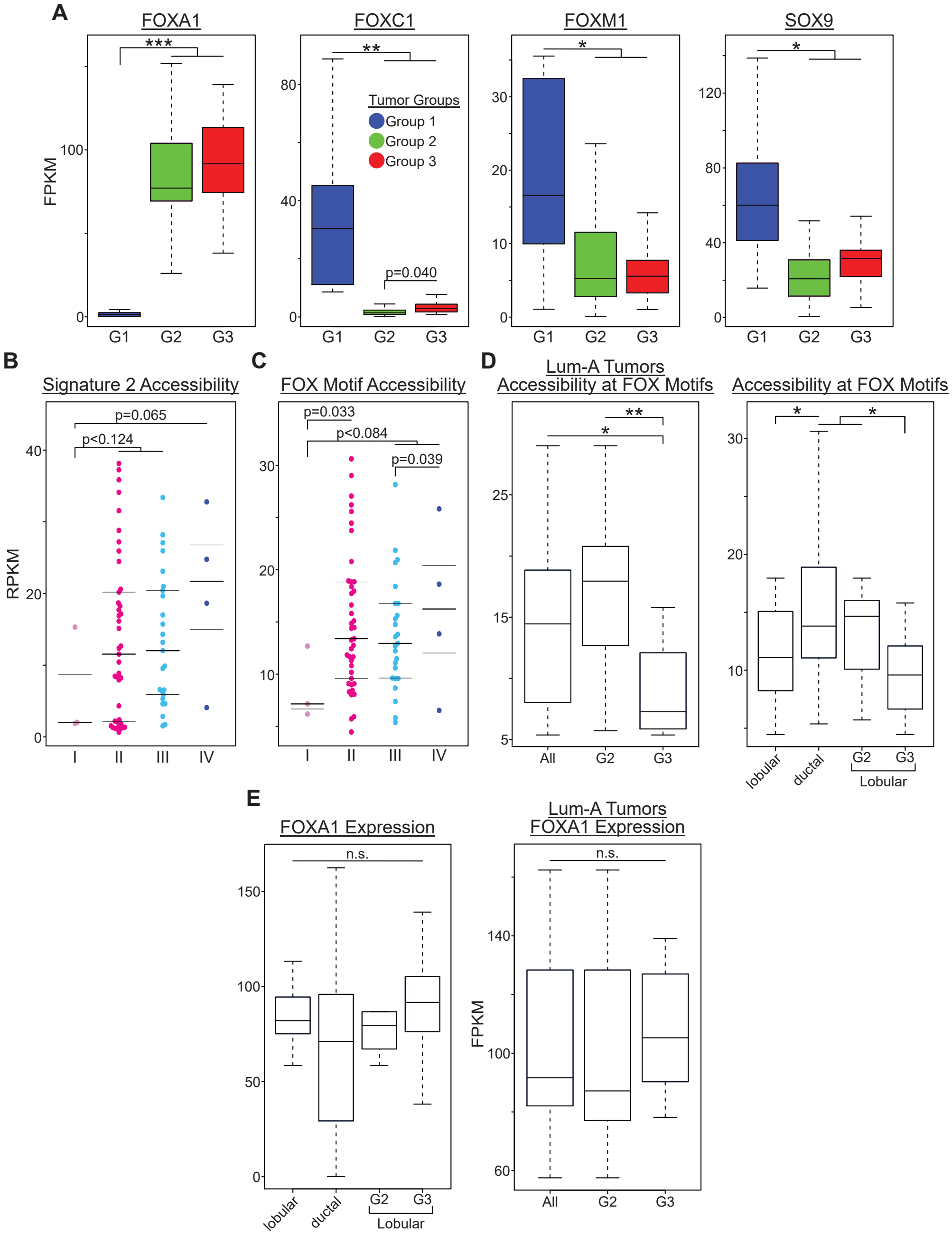


Figure 4 supplement:

A) Boxplots comparing expression across tumor groups of interesting SOX and FOX factors from figure 2G. B-C) Beeswarm plots comparing tumor's average accessibility in signature 2 regions (n=1314) (B) and tumor's average accessibility at FOX motifs within accessible peak regions (n=96280) (C) by tumor stage. D) Boxplots of average accessibility at FOX motifs in accessible peak regions of Luminal-A tumors separated into all (n=25), group 2 (n=17) and group 3 (n=8) (left) and of ductal (n=55) and lobular tumors separated into all (n=14), group 2 (n=8), and group 3 (n=6) (right). E) Boxplots of *FOXA1* expression for same tumors as panel D. P-values in A-E obtained with one-tailed parametric t-tests. * is p<0.01, ** is p<0.001, *** is p<0.0001.

**Supp. Fig. 5**

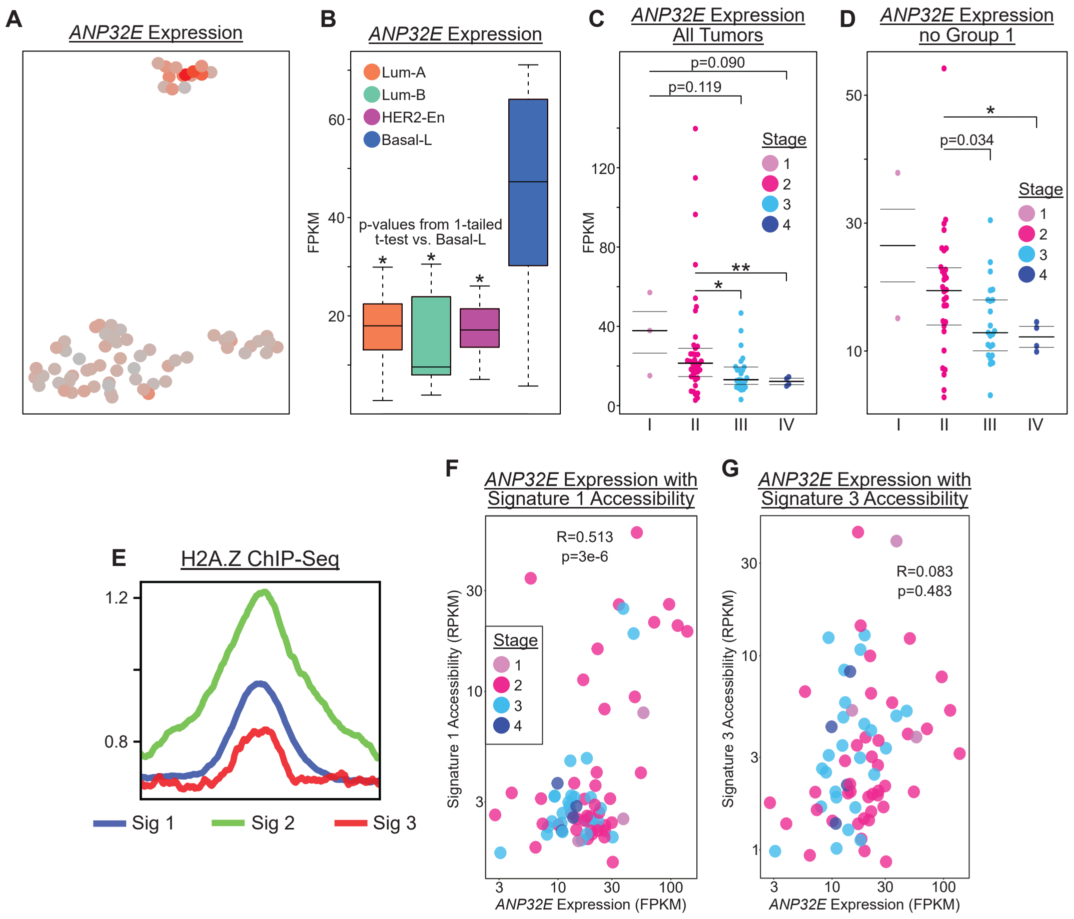


Figure 5 Supplement:

A) UMAP plot depicting *ANP32E* expression of tumors (n=74). B) Boxplot comparing *ANP32E* expression of tumors by PAM50 subtype. C-D) Beeswarm plots comparing *ANP32E* expression by tumor stage for all tumors with available stage data (n=73) (C) and for only group 2 and 3 tumors (n=59) (D). E) Profile plot showing binding of H2A.Z in MCF-7 cells within regions from signatures 1, 2 and 3. Data from ChIP-Seq of MCF-7 cells. F-G) Scatterplots showing the correlation of *ANP32E* expression with tumor's average accessibility in signature 1 (n=7026) (F) and signature 3 regions (n=527) (G). R denotes Pearson correlation coefficient; p-values from Pearson's product moment correlation coefficient. P-values in B-D obtained from one-tailed parametric t-tests. * is p<0.01, ** is p<0.001, *** is p<0.0001.

**Supp. Fig. 6**

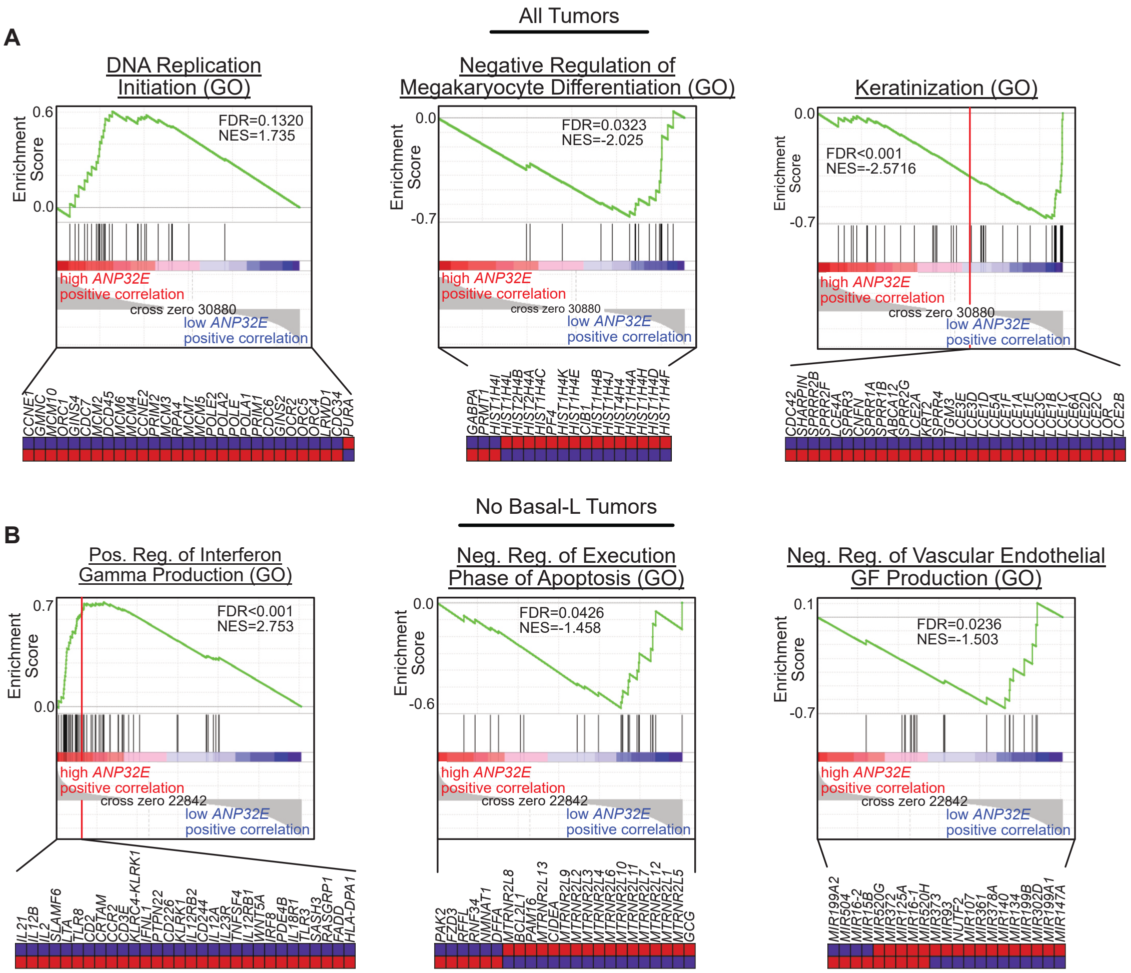


<u>Figure 6 Supplement</u>

A-B) GSEA plots depicting gene ontology associations with high and low *ANP32E* expression levels for all tumors with RNA-seq data from the TCGA-BRCA project (n=1222) (A) and for non-Basal-L tumors (n=1110) (B). FDR and NES values obtained within GSEA.

Supplemental Table 1 Legend:

HOMER identifies transcription factor motifs enriched within chromatin signature regions. Column 1 indicates the top 20 candidate transcription factor motifs for each signature group, along with reference numbers provided by HOMER databases specifying original motif discovery studies. Column 2 indicates p-values for significance of enrichment, as compared with background regions. Columns 3 and 4 indicate the percent of signature regions or background regions (respectively) which contain each candidate sequence motif.

Signature 1

| Motif Name | P-value | % in Signature | % in Background |
|---|---|---|---|
| Sox6(HMG)/(GSE32627) | 1.00E-71 | 29.42% | 20.41% |
| Sox9(HMG)/(GSE73225) | 1.00E-62 | 18.93% | 11.96% |
| Sox15(HMG)/(GSE62909) | 1.00E-56 | 22.13% | 14.96% |
| Sox2(HMG)/(GSE11431) | 1.00E-55 | 18.32% | 11.82% |
| Sox3(HMG)/(GSE33059) | 1.00E-52 | 31.80% | 23.74% |
| Sox10(HMG)/(GSE35132) | 1.00E-49 | 29.95% | 22.26% |
| Sox17(HMG)/(GSE61475) | 1.00E-44 | 15.00% | 9.68% |
| Sox4(HMG)/(GSE50066) | 1.00E-37 | 16.79% | 11.56% |
| CEBP(bZIP)/(GSE21512) | 1.00E-33 | 16.51% | 11.64% |
| RIN(MADS)/(GSE116581) | 1.00E-30 | 18.59% | 13.66% |
| Mef2c(MADS)/(GSE32465) | 1.00E-29 | 8.78% | 5.43% |
| Mef2b(MADS)/(GSE67450) | 1.00E-26 | 14.05% | 9.98% |
| NF1-halfsite(CTF)/(Unpublished) | 1.00E-24 | 34.81% | 29.08% |
| Mef2a(MADS)/(GSE21529) | 1.00E-19 | 7.91% | 5.28% |
| NFkB-p65(RHD)/(GSE19485) | 1.00E-18 | 7.80% | 5.26% |
| Pitx1(Homeobox)/(GSE38910) | 1.00E-16 | 53.36% | 48.32% |
| GRHL2(CP2)/(GSE46194) | 1.00E-15 | 13.83% | 10.74% |
| REM19(REM)/(GSE60143) | 1.00E-15 | 13.29% | 10.27% |
| HIC1(Zf)/(GSE99889) | 1.00E-15 | 29.22% | 25.01% |
| ARF2(ARF)/(GSE60143) | 1.00E-15 | 31.13% | 26.84% |

Signature 2

| Motif Name | P-value | % in Signature | % in Background |
|---|---|---|---|
| FOXM1(Forkhead)/(GSE72977) | 1.00E-98 | 67.88% | 39.01% |
| FOXA1(Forkhead)/(GSE27824) | 1.00E-83 | 70.32% | 43.75% |
| FOXA1(Forkhead)/(GSE26831) | 1.00E-82 | 66.29% | 39.88% |
| Fox:Ebox(Forkhead,bHLH)/(GSE47459) | 1.00E-73 | 52.82% | 28.81% |
| Foxa2(Forkhead)/(GSE25694) | 1.00E-70 | 55.02% | 31.18% |
| Foxa3(Forkhead)/(GSE77670) | 1.00E-53 | 36.83% | 18.63% |
| PHA-4(Forkhead)/(modEncode) | 1.00E-40 | 81.28% | 64.49% |
| FOXK1(Forkhead)/(GSE51673) | 1.00E-29 | 46.73% | 31.69% |
| Foxf1(Forkhead)/(GSE77951) | 1.00E-25 | 46.12% | 31.99% |
| FoxD3(forkhead)/(GSE106676) | 1.00E-25 | 39.27% | 25.83% |
| FoxL2(Forkhead)/(GSE60858) | 1.00E-23 | 42.85% | 29.55% |
| FOXK2(Forkhead)/(E-MTAB-2204) | 1.00E-18 | 27.93% | 17.87% |
| WRKY27(WRKY)/(GSE60143) | 1.00E-17 | 21.31% | 12.74% |
| Foxo3(Forkhead)/(E-MTAB-2701) | 1.00E-16 | 34.55% | 24.09% |
| FOXP1(Forkhead)/(GSE31006) | 1.00E-15 | 23.36% | 14.75% |
| WRKY28(WRKY)/(GSE60143) | 1.00E-15 | 26.33% | 17.35% |
| Hnf6b(Homeobox)/(GSE106305) | 1.00E-15 | 24.12% | 15.51% |
| WRKY29(WRKY)/(GSE60143) | 1.00E-14 | 22.83% | 14.82% |
| Foxo1(Forkhead)/(Fan_et_al.) | 1.00E-12 | 43.07% | 33.31% |
| COUP-TFII(NR)/(GSE46497) | 1.00E-12 | 20.78% | 13.52% |

Signature 3

| Motif Name | P-value | % in Signature | % in Background |
|---|---|---|---|

| | | | |
|---|---|---|---|
| CEBP(bZIP)/(GSE21512) | 1.00E-47 | 37.38% | 12.47% |
| HLF(bZIP)/(GSE69817) | 1.00E-33 | 33.59% | 13.05% |
| PPARa(NR),DR1/(GSE47954) | 1.00E-29 | 26.94% | 9.52% |
| PPARE(NR),DR1/(GSE13511) | 1.00E-24 | 24.10% | 8.90% |
| NFIL3(bZIP)/(Encode) | 1.00E-22 | 25.43% | 10.27% |
| CEBP:AP1(bZIP)/(GSE21512) | 1.00E-22 | 26.94% | 11.35% |
| Atf4(bZIP)/(GSE35681) | 1.00E-18 | 14.99% | 4.78% |
| RXR(NR),DR1/(GSE13511) | 1.00E-17 | 23.72% | 10.39% |
| Chop(bZIP)/(GSE35681) | 1.00E-17 | 12.52% | 3.59% |
| EBF2(EBF)/(GSE97114) | 1.00E-11 | 18.22% | 8.62% |
| NF1-halfsite(CTF)/(Unpublished) | 1.00E-11 | 42.31% | 28.46% |
| AARE(HLH)/ | 1.00E-07 | 5.69% | 1.74% |
| Erra(NR)/(GSE31477) | 1.00E-06 | 30.36% | 20.68% |
| EBF(EBF)/(GSE21978) | 1.00E-06 | 5.31% | 1.68% |
| COUP-TFII(NR)/(Encode) | 1.00E-06 | 20.68% | 12.84% |
| THRb(NR)/(GSE52613) | 1.00E-06 | 50.28% | 39.58% |
| Rbpj1(?)/(GSE47459) | 1.00E-04 | 22.96% | 15.80% |
| NLP7(RWPRK)/(GSE60143) | 1.00E-04 | 26.00% | 18.53% |
| LBD18(LOBAS2)/(GSE60143) | 1.00E-04 | 35.10% | 27.17% |
| EAR2(NR)/(Encode) | 1.00E-04 | 17.46% | 11.59% |
